# Broad Effects of Activation of Alcohol Dehydrogenase 1 on Healthspan Extension

**DOI:** 10.1101/2025.11.23.687054

**Authors:** Avery Sukienik, Samuel Berhanu, Abbas Ghaddar, Elizabeth MacPhail, Eyleen J. O’Rourke

## Abstract

Nutritional, genetic, and pharmacological interventions can extend lifespan; however, fewer have been shown to extend healthspan—the period of life free from chronic, debilitating diseases. In line with this, the molecular effectors that drive healthspan are even less understood than those responsible for lifespan extension. We recently reported that activation of Alcohol Dehydrogenase 1 (ADH-1) extends lifespan in yeast and *C. elegans*. In addition, *adh-1* is transcriptionally activated in yeast, worms, mice, and humans in response to caloric restriction—an intervention that extends not only lifespan but also healthspan. Therefore, we investigated whether activating *adh-1* could also extend healthspan. We demonstrate here that *adh-1* activation has broad and robust effects on health, including resistance to age-related obesity, delayed sarcopenia, and attenuated neurodegeneration. Mechanistically, ADH-1-driven healthspan extension is associated with improved proteostasis. These findings position ADH-1 as a promising target for future research aimed at promoting healthy aging.

## Introduction

Healthspan is the length of life spent free from serious illness or disability. Although human life expectancy has increased by approximately 30 years over the past century(Crimmins, 2015), healthspan extension has not kept pace—a phenomenon referred to as the *healthspan gap* (Rowe & Kahn, 1987). This gap is also observed in laboratory conditions following genetic, nutritional, and pharmacological interventions that extend lifespan. For instance, *Caenorhabditis elegans* carrying loss-of-function mutations in *clk-1*, a gene encoding a protein required for coenzyme Q (a.k.a. ubiquinone-9, a key component of the electron transport chain) biosynthesis, exhibit increased longevity but also severe developmental and motility defects that are partially diet-dependent(Branicky et al., 2000; Hihi et al., 2002). Even the best-characterized long-lived *C. elegans* mutant, *daf-2*—which encodes the worm’s sole insulin/IGF-1 receptor, shows pronounced developmental defects(Gems et al., 1998). Vertebrate longevity models also commonly display healthspan gaps. For example, mice carrying point mutations in codon 72 of the tumor suppressor p53 are long-lived(Migliaccio et al., 1999) but exhibit increased cancer susceptibility(Zhao et al., 2018). Similarly, methionine-restricted mice have extended lifespan but reduced bone density(Ables & Johnson, 2017), and the most studied mouse models of longevity, Ames and Dwarf(Bartke et al., 2001), show strong developmental and fertility defects(Kano, 2013). Thus, across diverse organisms, lifespan-extending interventions do not necessarily lead to proportional healthspan gains.

We recently reported that elevated Alcohol Dehydrogenase I (ADH-1) activity is necessary for lifespan extension driven by caloric restriction, as well as by mTOR and insulin signaling inhibition in *C. elegans* (Ghaddar et al., 2023). Importantly, constitutive activation of ADH-1 was also sufficient to extend lifespan, not only in worms but also in yeast. Furthermore, we found that calorically restricted mice, rats, pigs, Rhesus monkeys, and humans exhibit elevated *Adh1*/ADH1 expression (Ghaddar et al., 2023), suggesting evolutionary conservation of this response. In that study, and in the present one, we used three independent transgenic *C. elegans* lines that constitutively overexpress *adh-1* (hereafter referred to as ADH-1^OE^). These lines were validated to exhibit increased *adh-1* mRNA and protein expression and, critically, chronically elevated alcohol dehydrogenase activity (Ghaddar et al., 2023). The absence of obvious detrimental phenotypes in these long-lived animals (Ghaddar et al., 2023) supports the notion that ADH-1 activation may represent a viable strategy to promote longevity. However, whether ADH-1 activation also extends healthspan—delaying the onset or severity of age-associated functional decline—remains unknown.

Here, we demonstrate that the sole overexpression of ADH-1 (ADH-1^OE^) is sufficient to robustly extend healthspan in *C. elegans* feeding *ad libitum*. ADH-1^OE^ animals are resistant to both age-related and diet-induced obesity and exhibit reduced sarcopenia and neurodegeneration. Mechanistically, ADH-1^OE^ animals show transcriptional activation of a proteostatic program, accompanied by increased resistance to protein aggregation. These findings establish ADH-1 as a targetable driver of healthspan extension, supporting its potential as a therapeutic intervention against age-related decline.

## Results

### ADH-1^OE^ *C. elegans* are Resistant to Age-related and Diet-induced Obesity

Adiposity increases with age and is associated with the overall decline in health observed in aging animals (Bischof & Park, 2015; Palmer & Jensen, 2022; Palmer & Kirkland, 2016). Across organisms, life-extending interventions have diverse effects on adiposity, ranging in *C. elegans* from leanness under caloric restriction (Klapper et al., 2011) to obesity in long-lived notch (*glp-1*) and insulin receptor (*daf-2*) mutants (O’Rourke et al., 2009). To establish whether ADH-1 activity influences age-driven changes in adiposity, we first tested whether age-related obesity can be captured in *C. elegans* using the fat-specific dye Oil Red O (ORO) (Ke et al., 2018; Wählby et al., 2014a). For this, we stained young (day 1) and aged (day 10) wild-type animals. As anticipated, we observed higher ORO signal in aged worms (Fig. 1B), confirming that age-related fat accumulation can be effectively modeled and assessed in *C. elegans*.

**Figure 1.**
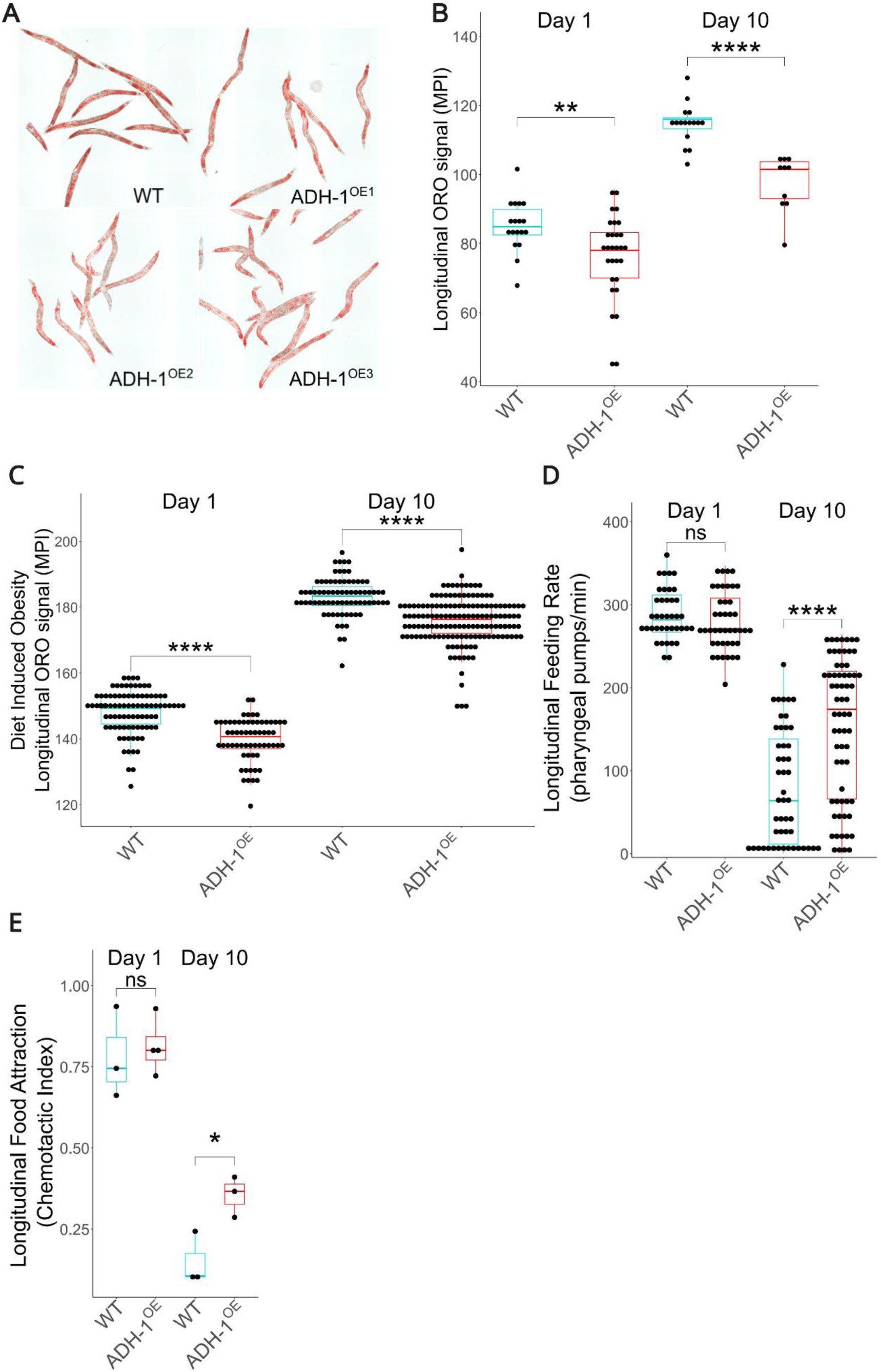
ADH-1 overexpression causes resistance to age-driven and diet-induced obesity without compromising feeding capacity. (A) Representative images of wild-type (WT) and ADH-1^OE^ *C. elegans* stained with the neutral-fat dye Oil Red O (ORO) at day 10 of adulthood (n=3 biological replicates). (B) Quantification of rORO intensity in WT and ADH-1^OE^ animals fed a standard *E. coli* OP50 diet at day 1 and day 10 of adulthood (n=3 biological replicates). (C) Quantification of ORO intensity in WT and ADH-1^OE^ animals fed an obesogenic High Fructose Diet (HFrD) (n=3 replicates, refer to DataS1C). (D) Quantification of pharyngeal pumping in WT and ADH-1^OE^ animals at day 1 and day 8 of adulthood (n=3 biological replicates). (E) Chemotactic index (CI) of WT and ADH-1^OE^ animals at day 1 and day 10 of adulthood (n=3 biological replicates, ≥100 worms per experimental replicate). Statistical testing was conducted using Welch’s t-test after outlier correction or Wilcoxon when populations were non-normal (Panels D and E). Error bars denote SEM. ns= not significant, *p<0.05, **p<0.01, ***p<0.001, ****p<0.0001, **MPI** = Mean Pixel Intensity.

*C. elegans* overexpressing ADH-1 (ADH-1^OE^) exhibit approximately 5-fold higher *adh-1* expression than wild-type animals (Ghaddar et al., 2023)–a level comparable to the induction observed under caloric restriction, a condition known to mobilize fat stores in *C. elegans* and other organisms (O’Rourke et al., 2009; Ghaddar et al., 2023). This parallel led us to test whether sustained ADH-1 activation could be sufficient to promote fat mobilization and counteract age-associated obesity. Indeed, ADH-1^OE^ worms markedly reduced lipid stores by day 10 of adulthood (Fig. 1A, B), suggesting improved metabolic health with age. Interestingly, even at day 1, ADH-1^OE^ worms were slightly leaner than wild-type controls (Fig. 1C, DataS1A, B).

To rule out the possibility that the observed reduction in Oil Red O (ORO) staining reflected altered cuticle permeability rather than true fat depletion, we tested staining across different incubation times. The rationale was that the cuticle becomes more permeable with age, and if ADH-1^OE^ animals remained more “youthful,” their cuticles might take longer to absorb the dye. However, we found that the lean phenotype of aged ADH-1^OE^ animals was consistent regardless of staining duration and was evident even after ORO saturation of ADH-1^OE^ animals had been reached (DataS1C), supporting a genuine reduction in fat content.

The experiments we describe were performed using the PHX2888 strain, one of three independently generated ADH-1^OE^ lines. In our previous work (Ghaddar et al., 2023), we characterized three independent transgenic lines overexpressing ADH-1 (GMW020-22). We found them to exhibit indistinguishable phenotypes across multiple biological scales, including *adh-1* expression levels, enzymatic activity, subcellular localization, and organismal phenotypes such as lifespan extension. Likewise, the lean phenotype was consistent across all three independent ADH-1^OE^ strains (DataS1D-F). We selected the GMW020 line—hereafter referred to simply as ADH-1^OE^—for the detailed healthspan analyses presented in this study, as it showed an intermediate degree of both leanness and lifespan extension.

Because aging also increases susceptibility to diet-induced metabolic dysregulation, we next tested whether ADH-1 activity could counter the obesogenic effects of fructose feeding in aged *C. elegans* (Ke et al., 2021). Consistent with this hypothesis, ADH-1^OE^ animals fed a fructose-enriched diet accumulated less fat than wild-type controls at both days 1 and 10 (Fig. 1C, DataS1G).

Because reduced fat stores in aged animals could reflect impaired feeding or abnormal food perception rather than improved metabolic health, we directly tested these possibilities at day 1 and 10. First, we found that young ADH-1^OE^ worms displayed normal contractility relative to age-matched wild-type; however, aged ADH-1^OE^ worms displayed better contractility than WT worms (Video 1, Video 2; Fig. 1D), indicating enhanced rather than diminished feeding capacity. We then assessed whether reduced adiposity might arise from defects in food detection or active avoidance. We found that young ADH-1^OE^ worms displayed normal response to food cues relative to WT worms; however, aged ADH-1^OE^ displayed better response to food cues relative to WT worms. (Fig. 1E). Together, these controls support the conclusion that reduced fat stores in aged ADH-1^OE^ animals reflect improved metabolic homeostasis, rather than feeding or sensory defects. The results also suggest better muscle function (pharynx) and locomotory and sensory capacity (chemotaxis) in ADH-1^OE^ *C. elegans*.

### ADH-1^OE^ *C. elegans* are Resistant to Oxidative Stress

Fats stored in lipid droplets, like those stained by ORO, can act as sinks for reactive oxygen species (ROS) that would otherwise cause broad-spectrum cellular damage (Bailey et al., 2015). For instance, insulin-insensitive *daf-2* mutant *C. elegans*—which have larger-than-wild-type fat stores—are more resistant to oxidative stress (Honda & Honda, 1999). Given that overexpressing ADH-1 reduces fat stores, we considered the possibility that this might compromise the animals’ resistance to oxidative stress.

To test whether reduced lipid buffering might increase ROS toxicity in ADH-1^OE^ animals, we exposed them to paraquat and measured survival, as previously described (Ishii et al., 1990; Mony et al., 2021; Senchuk et al., 2017). Surprisingly, ADH-1^OE^ animals exhibited enhanced survival compared to wild-type controls (Fig. 2A and B, DataS2 A-C), suggesting that fat depletion in these animals does not impair, but instead improves oxidative stress resilience.

**Figure 2.**
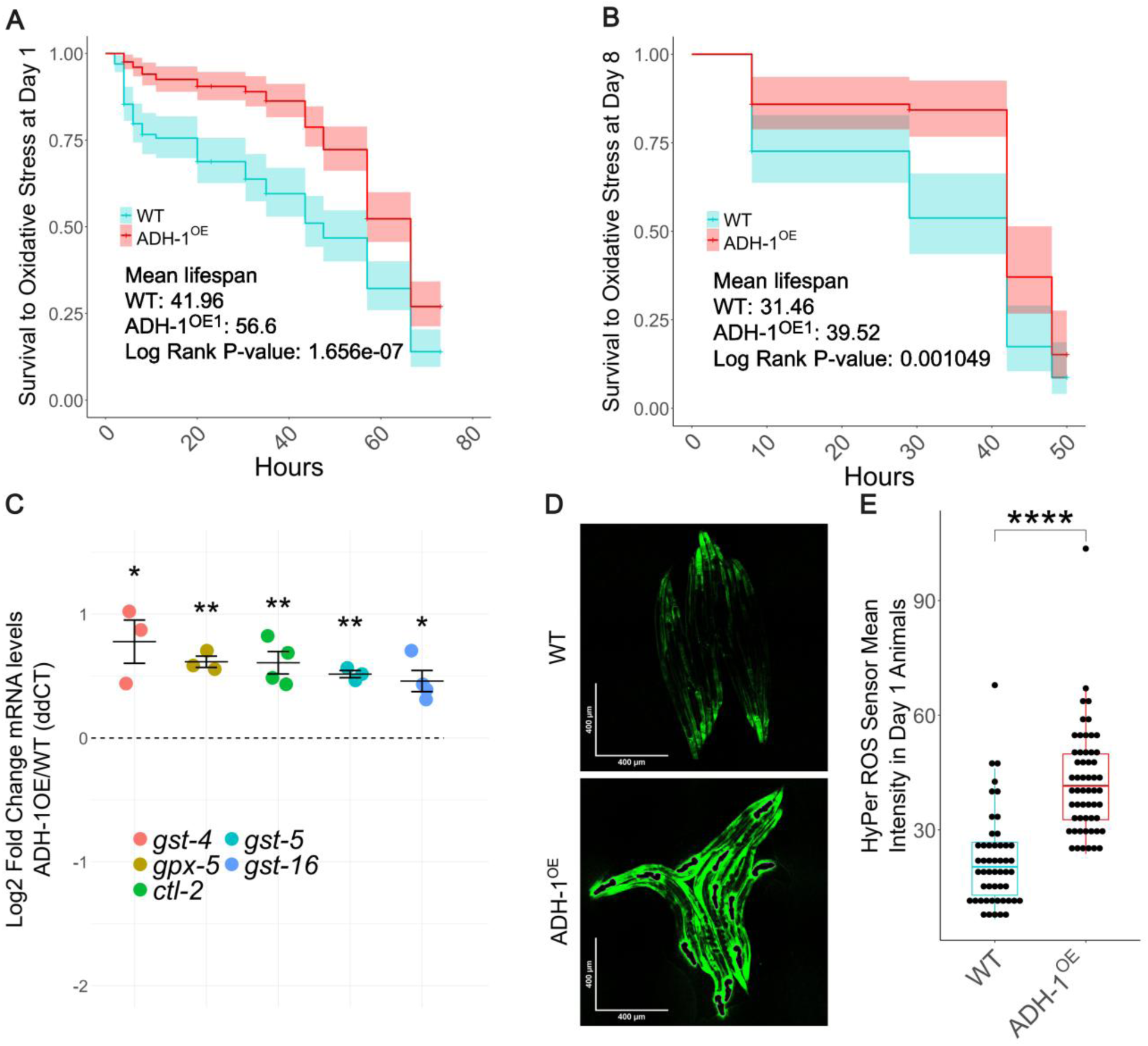
ADH-1 overexpression increases resistance to oxidative stress. (A) Day-1 ADH-1^OE^ *C. elegans* adults show enhanced survival to 100mM paraquat when compared to WT (representative of 2 biological replicates, see also DataS2A). (B) Day-8 ADH-1^OE^ *C. elegans* adults show enhanced survival to 100mM paraquat when compared to WT (representative of 3 biological replicates, see also DataS2B and C). (C) Day-1 ADH-1^OE^ adults show constitutive activation of an anti-oxidant transcriptional program (n=4 biological replicates, see also DataS2D), (D) ADH-1^OE^ animals show heightened H_2_O_2_ levels at day 1 of age as indicated by the HyPer::YFP H_2_O_2_ sensor, (E) Quantification of HyPer::YFP signal showing ADH-1^OE^ animals have increased ROS levels when compared to WT (n=3 biological replicates). Statistical testing was conducted using Welch’s t-test after outlier correction or Wilcoxon when populations were non-normal (Panel E). Statistical testing of survival conducted by Kaplan-Meier Log-Rank Fold test. Error bars denote SEM. ns= not significant, *p<0.05, **p<0.01, ***p<0.001, ****p<0.0001.

Consistent with this enhanced resistance, ADH-1^OE^ worms exhibit constitutively elevated expression of antioxidant genes, including *ctl-2, gpx-5, gst-4, gst-5, and gst-16* (Fig. 2C) (Back et al., 2012a; Hu et al., 2018). Notably, *ctl-2* has been directly implicated in protection from ROS-induced damage(Musa et al., 2022; Petriv & Rachubinski, 2004). Additional redox-responsive genes such as *ctl-3* and *sod-1* also trended toward upregulation in ADH-1^OE^ worms (DataS2D), consistent with a broader activation of antioxidant defenses (Blackwell et al., 2015).

Mild elevations in ROS levels have been documented in other long-lived *C. elegans* mutants, such as *clk-1* and *sod-2*, and in some cases are essential for their extended lifespan(Raamsdonk & Hekimi, 2009; Yang & Hekimi, 2010b). Similarly, low-dose oxidant exposure (e.g., juglone or paraquat) has been shown to increase lifespan via hormesis—whereby transient oxidative stress triggers protective responses that increase long-term survival(Heidler et al., 2010; Lee et al., 2010; Yang & Hekimi, 2010a). This model suggests that improved survival following mild oxidative stress may arise not from targeted adaptation to each specific insult that an animal could encounter throughout its lifespan, but from a generalized enhancement of stress resilience. Such broad-spectrum protection could defend against diverse ROS-linked threats encountered throughout life, including fatty acid oxidation, pathogen attack and its associated immune responses(Back et al., 2012b; Ristow & Schmeisser, 2011; Schulz et al., 2007).

To determine whether ADH-1 ^OE^ animals exhibit elevated endogenous ROS that might trigger such a hormetic response, we measured hydrogen peroxide (H₂O₂) levels using the HyPer biosensor—a genetically encoded, ratiometric probe combining a circularly permuted YFP with the H₂O₂-sensitive regulatory domain of *E. coli* OxyR (Belousov et al., 2006). ADH-1^OE^ day 1 adults displayed significantly elevated H₂O₂ levels compared to wild-type animals (Fig. 2D, E), consistent with a state of mild endogenous oxidative stress. In line with this increase, ADH-1^OE^ animals survived paraquat exposure better than controls and showed upregulation of multiple antioxidant defense genes. These findings support a model in which enhanced ADH-1 activity initiates a preconditioning response that boosts redox detoxification capacity. Thus, rather than sensitizing animals to oxidative damage, ADH-1 overexpression appears to enhance oxidative stress resistance, potentially through hormetic activation of stress response pathways.

### ADH-1^OE^ Enhances ER Stress Response

Lean phenotypes and resistance to oxidative stress have been associated with enhanced endoplasmic reticulum (ER) function and improved protein homeostasis (proteostasis) in multiple model systems, including *C. elegans* and mammals(Ben-Zvi et al., 2009; Henis-Korenblit et al., 2010; Taylor & Dillin, 2013a). A growing body of evidence suggests that low-level UPR activation can enhance the cellular capacity to manage misfolded proteins and oxidative stress, promoting metabolic resilience and extending organismal healthspan (Calfon et al., 2002a; Hetz & Papa, 2018; Wang & Kaufman, 2016). In this context, the UPR acts not as a sign of dysfunction but as an adaptive, preemptive defense mechanism.

To explore whether the improved metabolic and stress resilience of ADH-1^OE^ animals might stem from enhanced ER homeostasis, we examined the expression of *hsp-4*, a key component of the unfolded protein response and the *C. elegans* ortholog of mammalian BiP/GRP78. Using a *hsp-4*P::GFP reporter, we found that *hsp-4* expression is modestly but consistently elevated in ADH-1^OE^ worms under non-stressed conditions, particularly in the intestine and tail regions—tissues central to metabolic regulation (Fig. 3A and B).

**Figure 3.**
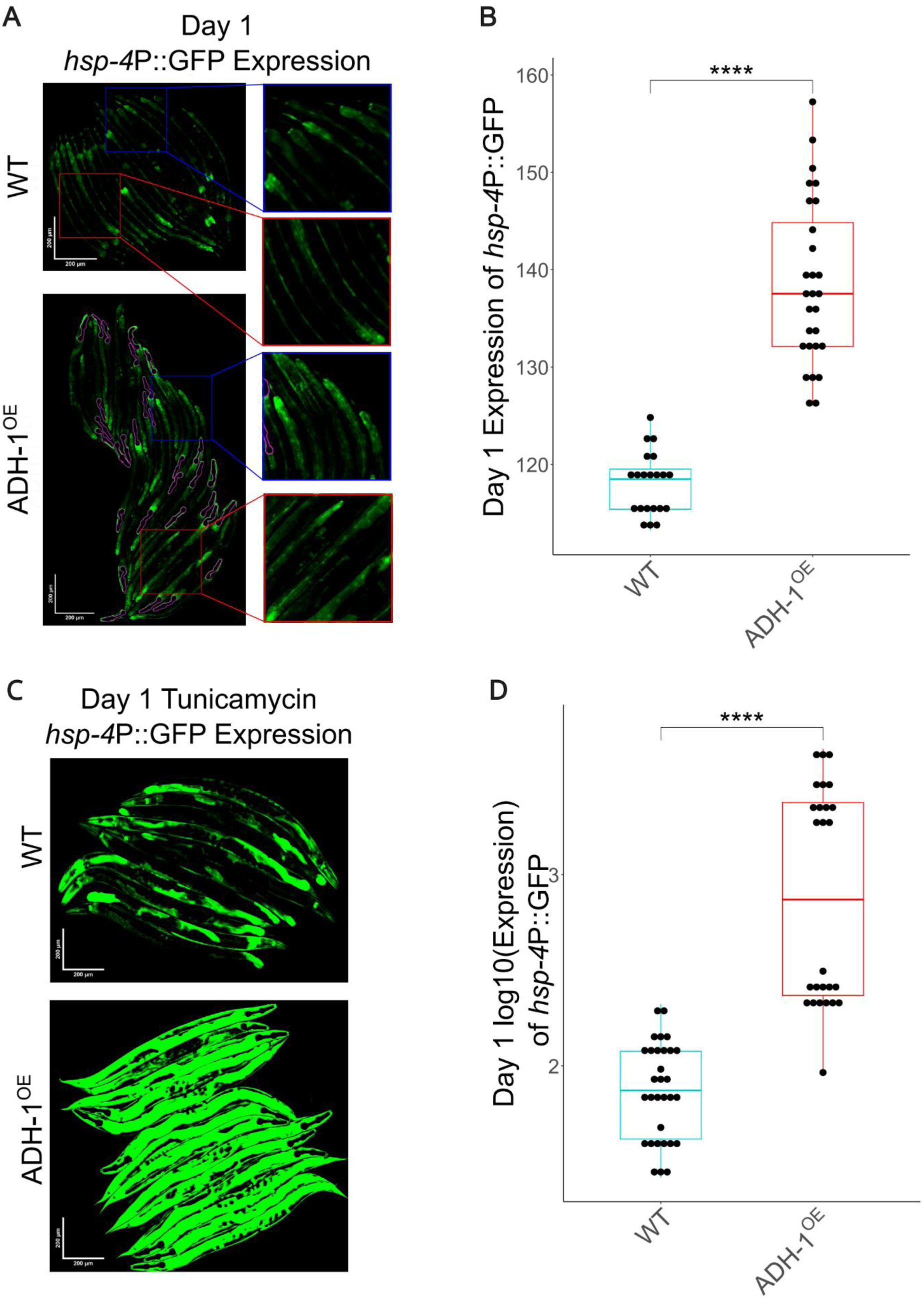
Activating ADH-1 causes mild activation of ER stress response. (A) Representative images showing that ADH-1^OE^ animals show enhanced expression of the gene encoding the ER-specific chaperone HSP-4 in the intestine–mainly in the anterior and posterior intestine, which accumulate most fat stores in *C. elegans*. (B) Quantification of *hsp-4*P::GFP intensity in unstressed wild-type and ADH-1^OE^ *C. elegans* as shown in panel A (n=3, see also DataS3A-B). (C) Quantification of *hsp-4*P::GFP intensity in wild-type *C. elegans* stressed with 5µg/µl of Tunicamycin, a treatment that induces pathological ER stress (n=2). Error bars denote SEM. Statistical testing was conducted using Welch’s t-test after outlier correction or Wilcoxon when populations were non-normal (Panels B and D). ns= not significant, *p<0.05, **p<0.01, ***p<0.001, ****p<0.0001.

Importantly, while *hsp-4*P::GFP levels are elevated at baseline in ADH-1^OE^ animals, this increase is far more modest than the strong induction observed following exposure to 5 µg/mL tunicamycin, a well-established ER stressor(Calfon et al., 2002b) (Fig. 3C and D). This distinction suggests that the elevation in *hsp-4* expression in ADH-1^OE^ animals likely reflects a mild, hormetic activation of the UPR rather than a pathological stress response. In turn, this preconditioning may enhance the animal’s capacity to cope with cumulative ER stress, potentially contributing to its improved metabolic health and oxidative stress resistance.

### ADH-1^OE^ Delays Age-dependent Loss of Locomotory Capacity

The improved pharyngeal pumping observed in aged *C. elegans* overexpressing ADH-1 (Fig. 1D) suggests that ADH-1^OE^ enhances the function of aging striated autonomic muscles. We next examined whether skeletal muscle integrity is similarly preserved in these animals. To assess this, we measured locomotory function using the thrashing assay, which quantifies the number of body bends per second (bbps) as worms respond to liquid immersion. This behavior declines with age and is a widely used proxy for neuromuscular health in *C. elegans*(Dillin et al., 2002; Glenn et al., 2004; Herndon et al., 2002; Hosono, 1978; Hosono et al., 1980; Hsu et al., 2009; C. Huang et al., 2004; Johnson et al., 1988; Liu et al., 2013). At day 1 of adulthood, locomotory activity was similar between wild-type and ADH-1^OE^ animals (Fig. 4A). However, by day 12, ADH-1^OE^ worms maintained significantly higher bbps compared to age-matched controls (Fig. 4A; DataS4A-B), indicating that the locomotory benefits of ADH-1^OE^ emerge with age.

**Figure 4.**
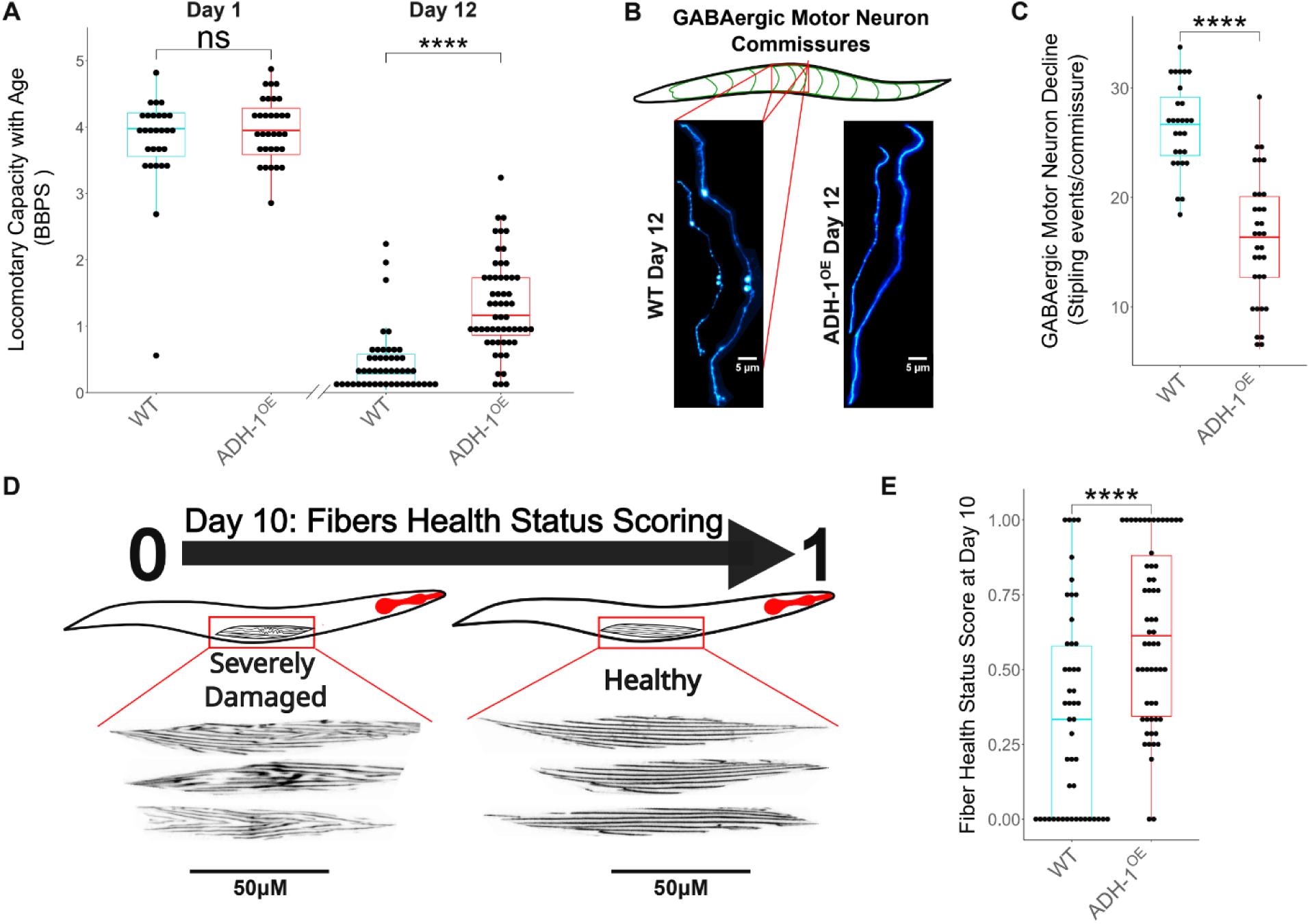
ADH-1 Overexpression Preserves Motor Neuron and Skeletal Muscle Structure During Aging. (A) Quantification of locomotion capacity through the thrashing assay shows no baseline advantage of ADH-1^OE^ animals at day 1 (n = 3 biological replicates), but at day 12, ADH-1^OE^ animals exhibit more BBPS than WT animals, indicative of improved locomotory capacity relative to same-age WT *C. elegans* (representative of 3 biological replicates; see also DataS4). (B) Representative images of the axonal commissures of GABAergic motor neurons in WT (left) and ADH-1^OE^ (right) animals at day 12 (n = 2 biological replicates). (C) Quantification of GABAergic motor neurodegeneration showing that ADH-1^OE^ animals have less axonal stippling events per commissure than same-age WT worms (n = 2 biological replicates). (D) Schematic showing the muscle health scoring system used to assess sarcopenia status. (E) At day 10, the muscle health scores of ADH-1^OE^ animals are significantly higher than those of WT (n = 2 biological replicates). Statistical testing was conducted using Welch’s t-test after outlier correction, or Wilcoxon test when populations were non-normal (Panel A and F). ns = not significant; *p < 0.05; **p < 0.01; ***p < 0.001; ****p < 0.0001.

Age-dependent locomotory decline is typically accompanied by progressive deterioration of skeletal muscle fibers and motor neurons (Chow et al., 2006; Cruz-Jentoft et al., 2019; Dhondt et al., 2021; Giunti et al., 2024; X. Huang et al., 2002; Pan et al., 2011; Tank et al., 2011). Thus, the improved locomotory activity observed in aged ADH-1^OE^ worms suggests that elevated alcohol dehydrogenase activity would help preserve motor neuron and/or skeletal muscle function.

GABAergic motor neurons are particularly prone to early age-dependent decline, experiencing functional decline before the onset of sarcopenia (Glenn et al., 2004; Herndon et al., 2002). This decline is characterized most prominently by late-life structural abnormalities in the commissures—the axons that project from the ventral and dorsal cord neurons of *C. elegans* GABAergic motor neurons (Fig. 4C). Common abnormalities include bulges, beading, and fasciculation(Pan et al., 2011; Tank et al., 2011), collectively referred to as “stippling”, which serve as markers of overall neuronal health(Giunti et al., 2024). To determine whether the enhanced locomotory function in ADH-1^OE^ worms is associated with improved motor neuron maintenance, we examined commissure structure in aging worms(Gjorgjieva et al., 2014). In line with improved motor neuron maintenance, 12-day-old ADH-1^OE^ worms exhibited less commissure stippling than their wild-type counterparts (Fig. 4B & C). Notably, while WT control animals showed examples of extreme neurodegeneration (DataS4C), ADH-1^OE^ animals were not noted to have any such cases at day 12.

Having established that ADH-1^OE^ preserves motor neuron integrity with age, we next asked whether skeletal muscle structure might also be protected. Age-related locomotory decline is strongly associated with sarcopenia—the progressive degeneration of body wall muscles—which manifests cytologically as disorganized and degraded muscle fibers (Cruz-Jentoft et al., 2019, 2019; Dhondt et al., 2021). To assess whether the improved locomotory capacity in ADH-1^OE^ worms was also associated with preserved muscle architecture, we used a strain expressing a GFP::MYO-3 fusion protein(Campagnola et al., 2002), which allows visualization of the body wall muscle sarcomeres through green fluorescence. Sarcopenia was assessed by binary scoring of GFP::MYO-3 in the midbody: worms with well-structured sarcomeres received a score of 1, while those with severe sarcomere disorganization received a score of 0 (Fig. 4D, E). As expected, muscle fibers in both wild-type and ADH-1^OE^ worms exhibited parallel, well-organized sarcomere layers—most scoring as grade 1—in young day 1 adults (Fig. 4D, E). While sarcomere deterioration occurred with age in both genotypes, 10-day-old ADH-1^OE^ worms showed significantly better preservation of striated muscle structure (Fig. 4F). Interestingly, the proportion of intermediately damaged fibers was similar between groups (DataS4E, H), suggesting that muscle aging may proceed in discrete steps rather than as a gradual decline. This pattern is consistent with an age-dependent increase in severely damaged fibers in wild-type worms, a trajectory from which ADH-1^OE^ animals appear partially protected (DataS4D–I).

Together, these results suggest that the improved locomotory function observed in aged ADH-1^OE^ worms stems from the combined structural preservation of both GABAergic motor neurons and body wall muscles.

### ADH-1^OE^ Improves Muscle Proteostasis and Late-Age Locomotion

Impaired proteostasis is a fundamental driver of age-related decline in muscle structure and function across species, contributing to sarcopenia and reduced physical performance. In *C. elegans*, as in mammals, the progressive accumulation of misfolded and aggregated proteins in muscle cells underlies functional deterioration with age.(Hipp et al., 2019; Labbadia & Morimoto, 2014; Meller & Shalgi, 2021). Given that ADH-1 overexpression preserves muscle integrity and locomotory performance during aging, we tested whether it also improves muscle proteostasis. To address this, we employed two well-established *C. elegans* models of muscle-specific proteotoxic stress.

Most proteins fold optimally within a narrow temperature range, and deviations from this range activate the unfolded protein response (UPR) and recruit chaperones to assist in protein refolding. Thus, one approach to challenging the global proteostasis network is to expose animals to heat stress. Chronic exposure to 27°C is sufficient to induce proteostatic stress in *C. elegans*, providing a straightforward assay for the organism’s capacity to maintain protein homeostasis as it ages. Consistent with improved proteostasis, ADH-1^OE^ animals survived longer than wild-type controls under chronic heat stress (Fig. 5A, DataS5A), indicating enhanced resilience to protein misfolding stress.

**Figure 5.**
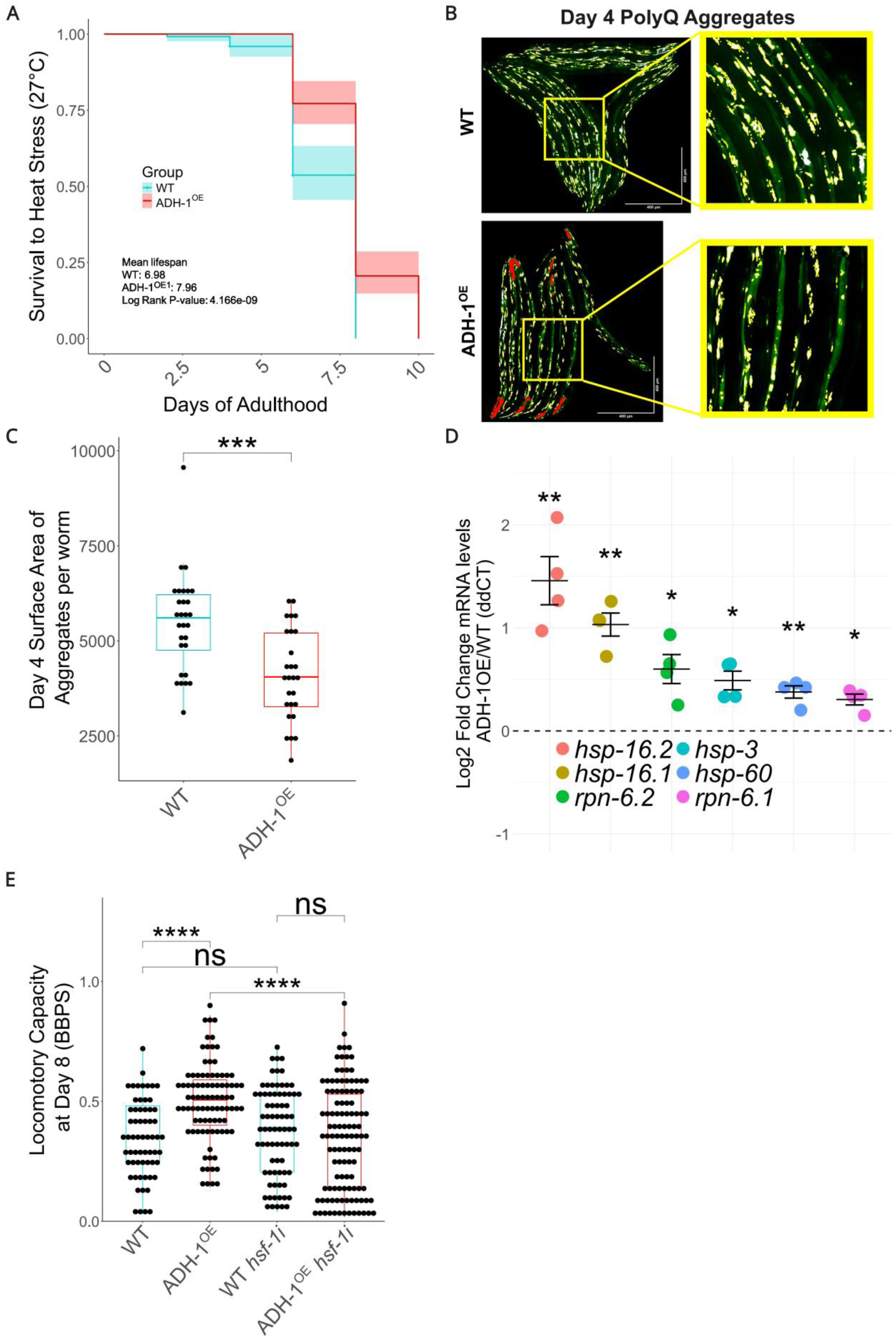
ADH-1 overexpression results in resistance to proteostatic stress. (A) Representative survival curve showing that ADH-1^OE^ animals better survive chronic heat stress (n=2 replicates, see DataS5A) (B) Representative images of day-4 animals expressing polyQ::YFP (depicted green) with masks highlighting the thresholded aggregates (depicted yellow to white according to their increasing signal intensity) (C) Quantification of total surface area of aggregates per animal shows that ADH-1^OE^ animals have less polyQ::YFP aggregation than same-age WT worms (n=3 biological replicates), (D) ADH-1^OE^ animals show increased expression of key proteostatic genes (n=4 biological replicates), (E) *hsf-1i* suppresses ADH-1^OE^ dependent locomotory capacity improvements at day 8 (n=2 biological replicate). Statistical testing conducted by Welch’s t-test after outlier correction or by Wilcoxon when populations were non-normal (Panel E). Error bars denote SEM. ns= not significant, *p<0.05, **p<0.01, ***p<0.001, ****p<0.0001.

While heat stress resistance is a widely used proxy for organismal proteostasis, it reflects the integrated response of multiple cellular pathways and stress axes, making it an informative but indirect readout of proteostatic capacity. To directly assess muscle-specific proteostasis, we next used a well-established model of muscle proteotoxicity in *C. elegans*: the AM140 strain, which carries the integrated transgene *rmIs132* (*unc-54p::Q35::YFP*). This construct expresses a polyglutamine (polyQ) stretch fused to YFP under control of the *unc-54* muscle-specific promoter, leading to age-dependent aggregation of the fusion protein in body wall muscle cells. In young AM140 animals, the YFP signal is diffuse and outlines the muscle fibers, producing a smooth contour along the body wall. As the animals age, the YFP signal becomes increasingly punctate due to protein aggregation, and the fiber-defined contour is lost—reflecting a decline in muscle proteostasis. (Ellis et al., 2023; Morley et al., 2002).

To test whether ADH-1 overexpression enhances muscle proteostasis, we crossed the *Q35::YFP* transgene into the ADH-1^OE^ background. Longitudinal analysis revealed that ADH-1^OE^ animals exhibited longer and more continuous stretches of diffuse YFP signal compared to wild-type animals, beginning as early as day 4 of adulthood (Fig. 5B-C). This observation suggests reduced aggregation and supports the conclusion that ADH-1^OE^ extends the period during which muscle proteostasis is maintained.

To investigate the molecular basis of this enhanced proteostasis, we measured the expression of several heat shock protein encoding genes (*hsp*), which are critical components of the proteostatic network. Quantitative PCR analysis showed that ADH-1^OE^ animals exhibit elevated expression of *hsp-3*, *rpn-6.1*, *rpn-6.2*, *hsp-16.1*, *hsp-16.2*, and *hsp-60* (Fig. 5D).

To test whether transcriptional activation of heat shock proteins plays a role in the locomotory benefits of ADH-1^OE^, we knocked down *hsf-1*, the master regulator of the heat shock response (Hajdu-Cronin et al., 2004; Morley & Morimoto, 2004). Although *hsf-1* RNAi had no effect on locomotion in day 1 adults (DataS5B), it suppressed the improved locomotory capacity of aged ADH-1^OE^ animals (Fig. 5F), suggesting that *hsf-1*-dependent transcriptional programs are required for the maintenance of locomotion in these animals.

Together, these findings suggest that ADH-1^OE^ enhances muscle proteostasis at multiple levels, including both tissue-specific reductions in protein aggregation and systemic improvements in heat stress resilience, likely mediated through *hsf-1*-dependent upregulation of HSPs. These results support a model in which ADH-1^OE^ extends healthspan not only by improving metabolic homeostasis, but also by maintaining proteostatic capacity in aging muscle through the activation of *hsf-1*–driven stress

### ADH-1 May Act Through a Non-Cell-Autonomous Mechanism

Our findings indicate that overexpression of *adh-1* preserves the function of multiple tissues in aging *C. elegans*, including GABAergic motor neurons (Fig. 3D). However, confocal imaging of overexpressing ADH-1 tagged to wrmScarlet (Fig. 6B) reveals a more restricted expression pattern for this enzyme. Specifically, ADH-1 is predominantly expressed in body wall muscle, pharyngeal muscle, and the intestine—but not in GABAergic motor neurons (Fig. 6C–F). Similarly, we observed no detectable ADH-1 expression in olfactory neurons such as AWB cells (Fig. 6D), which—although not directly examined here—are likely better maintained in ADH-1^OE^ animals, based on their improved chemotactic performance (Fig. 1E).

**Figure 6.**
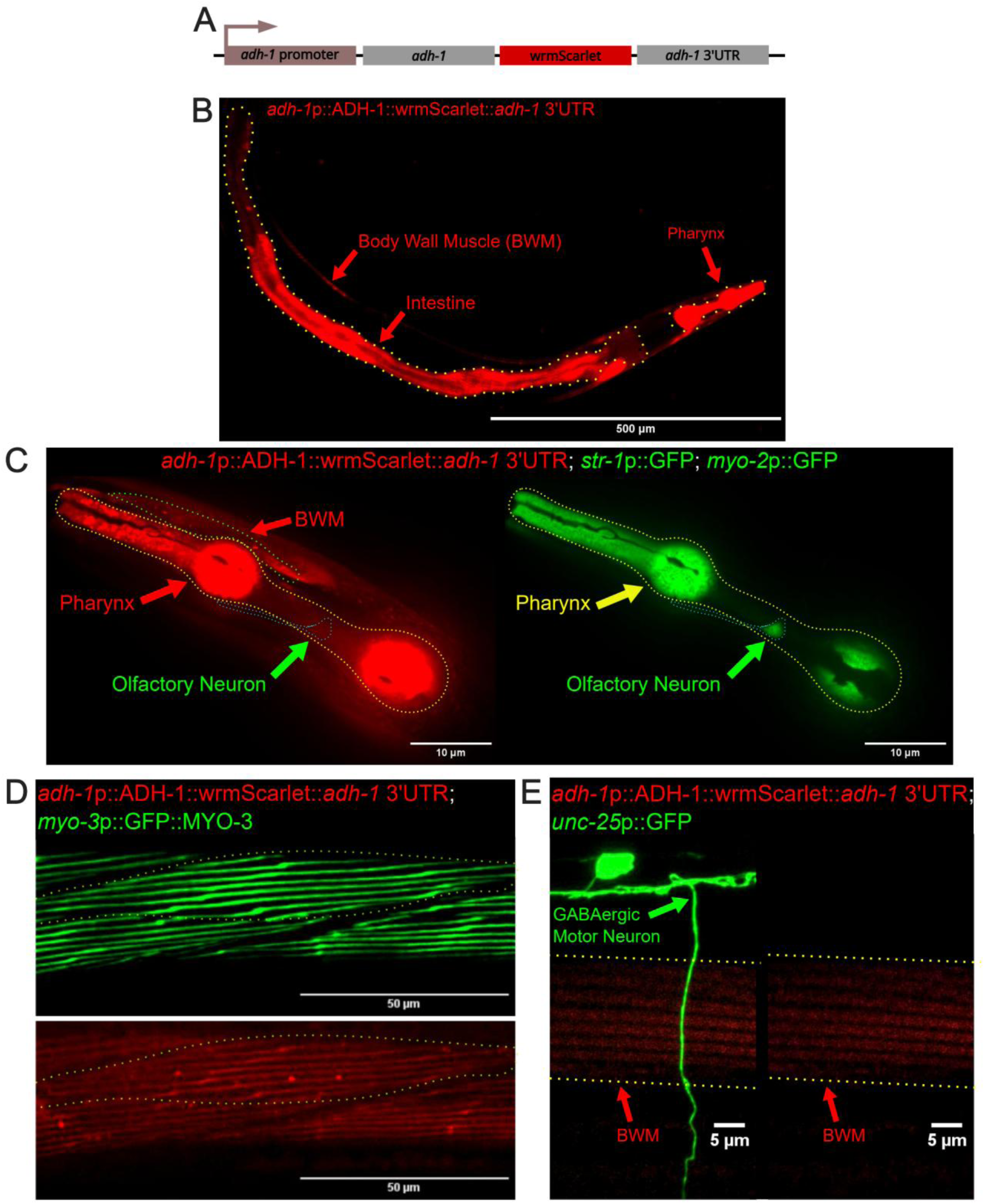
ADH-1 may act through a non-cell autonomous mechanism. (A) Schematic of the pertinent sequence used to generate ADH-1^OE^ animals (*adh-1*p::ADH-1::wrmScarlet*::adh-1* 3’UTR), (B) ADH-1 is strongly expressed in the intestine (outlined in yellow) and the pharynx, (C) ADH-1 is strongly expressed in the pharynx (yellow arrow) but not in AWB neurons, as indicated by lack of colocalization with *str-1*p::GFP (green arrow), (D) ADH-1 is expressed in body wall muscle (BWM), as defined by colocalization with *myo-3*p::GFP, (E) ADH-1 is not detectable in GABAergic motor neurons, as indicated by lack of colocalization with *unc-25*p::GFP. Muscle expression of *adh-1*::wrmScarlet is outlined in yellow for comparison.

Together, these findings suggest that the beneficial effects of ADH-1 overexpression extend beyond its sites of expression, supporting a cell non-autonomous mechanism of action. While the precise inter-tissue signals mediating these effects remain to be elucidated, our results establish that ADH-1 activation enhances healthspan across multiple systems in *C. elegans*. This observation sets the stage for future studies aimed at identifying systemic signals—such as metabolites, redox-active molecules, myokines, or other intercellular messengers—that may underlie the broad physiological improvements associated with ADH-1 activity.

## Discussion

Many established geroprotective interventions extend lifespan, but they often do so unevenly across organ systems, sometimes improving select features of healthspan while neglecting or impairing others. For example, loss-of-function in *clk-1* (CoQ7) extends lifespan but causes delayed development and reduced locomotion(Cristina et al., 2009). By contrast, ADH-1 overexpression yields coordinated health benefits across distinct physiological domains—including proteostasis, stress resilience, locomotion, neuronal maintenance, and leanness. This breadth of benefit invites consideration of how aging processes across tissues might be modulated by a common upstream factor.

At a mechanistic level, our findings suggest that ADH-1 promotes healthspan through metabolic rewiring that subtly alters cellular redox balance (Fig. 7).

**Figure 7.**
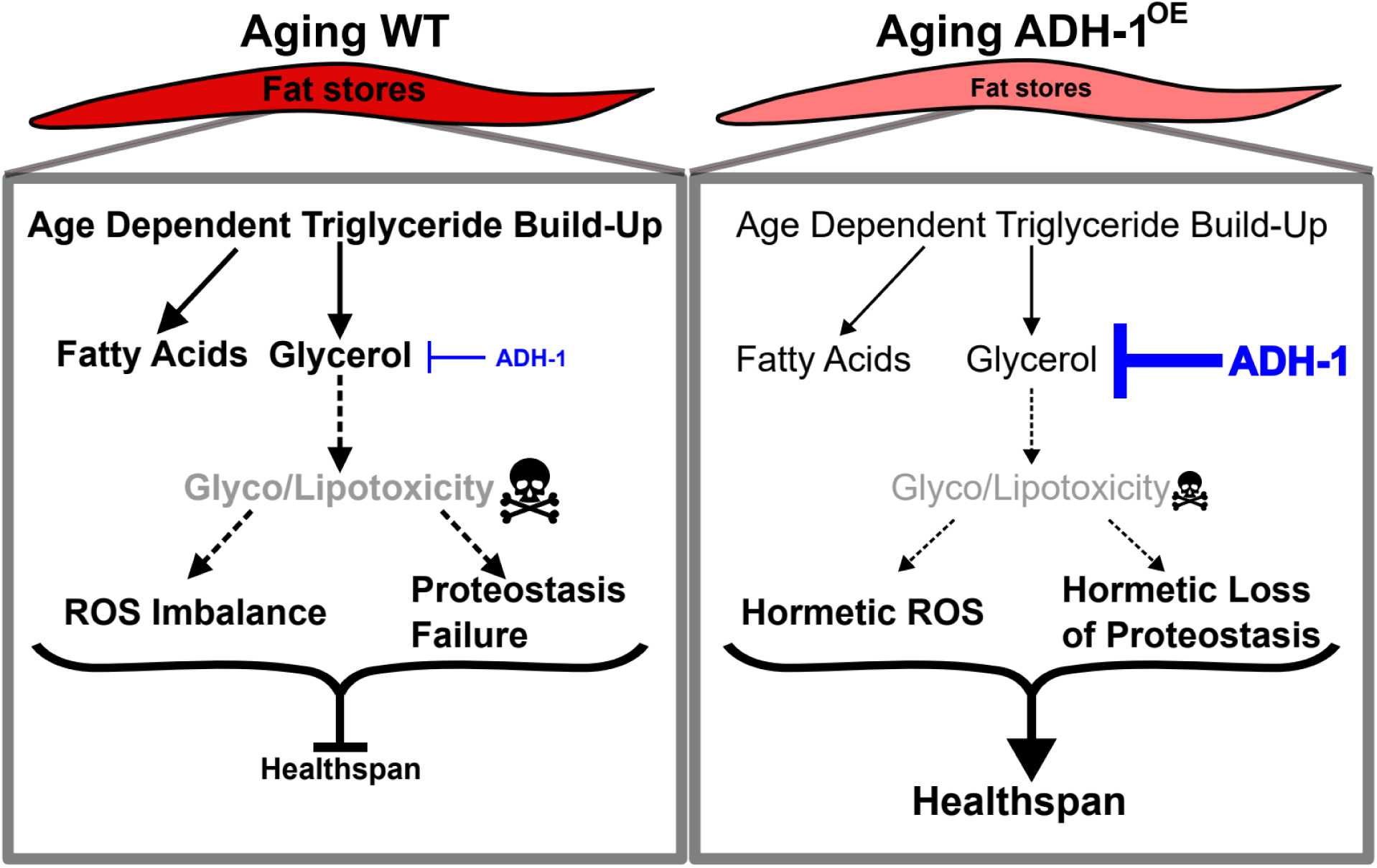
ADH-1 limits age-dependent glycerol build-up and tunes ROS and proteostasis to extend healthspan. Schematic model comparing aging wild-type (WT, left) and ADH-1–overexpressing (ADH-1^OE^, right) animals under nutrient-rich conditions. In aging WT, age-dependent triglyceride accumulation leads to glycerol build-up, which favors glyco-/lipotoxic stress (e.g., advanced glycation and lipid peroxidation). This stress drives ROS out of the hormetic range and contributes to proteostasis failure (inadequate UPR/HSR activity), resulting in only standard healthspan. By contrast, in ADH -1^OE^ animals, increased ADH-1 activity limits fat accumulation and, based on prior work, glycerol build-up, thereby reducing glyco-/lipotoxic stress. As a result, ROS and proteostasis mechanisms (e.g., chaperones) are maintained within a hormetic window, preserving tissue function and extending healthspan. Solid arrows and black font indicate relationships supported by data in this and in previous (Ghaddar et al., 2023) work, whereas dashed arrows and grey font depict proposed mechanistic links.

The modest but consistent increase in reactive oxygen species (ROS) observed in ADH-1^OE^ animals is reminiscent of mitohormesis, wherein low-level oxidative stress activates protective transcriptional programs that confer stress resistance and longevity(Ristow & Zarse, 2010; Yun & Finkel, 2014). Such hormetic ROS signaling has been implicated in longevity-promoting effects of metformin (De Haes et al., 2014), mitochondrial uncoupling (Cristina et al., 2009), and proline catabolism (Zarse et al., 2012).

Given ADH-1’s biochemical function in alcohol oxidation and its impact on NAD⁺/NADH balance, its overexpression may stimulate NAD⁺ cycling and enhance mitochondrial metabolism via β-oxidation and glycerol catabolism—both of which can influence ROS production and systemic energy homeostasis. Indeed, here we show that ADH-1 promotes leanness under nutrient-rich conditions (Ghaddar & Mony et al., 2023), positioning it as a modulator of intracellular flux and metabolic flexibility.

The redox shift caused by overexpressing ADH-1 appears to prime cytoprotective pathways. ADH-1^OE^ animals exhibit hallmarks of enhanced proteostasis, including upregulation of multiple hsp genes and activation of the ER unfolded protein response (UPR), as shown by both transcriptomic analyses and *hsp-4*p::GFP reporter expression. While the UPR is classically associated with cellular stress, moderate activation enhances protein folding capacity and extends lifespan in *C. elegans* and other models (Taylor & Dillin, 2013b). Disruptions in protein glycosylation or folding, potentially resulting from glucose limitation via glycerol diversion, could underlie the ER stress observed in ADH - 1^OE^ animals(Cruz-Rodríguez et al., 2025). ROS can also engage UPR sensors (e.g., IRE-1 and ATF6) through redox-sensitive modifications (Cao & Kaufman, 2014; Malhotra & Kaufman, 2007), providing a molecular basis for redox–UPR coupling. Importantly, our genetic data indicate that such coupling is not uniformly required for all ADH-1 benefits: while UPR priming may contribute to proteostasis and heat-stress resilience, RNAi against UPR branches did not eliminate the leanness phenotype (with only partial trends), suggesting that metabolic outcomes can be decoupled from IRE-1/ATF-6–dependent signaling in this context.

Muscle function is another critical determinant of healthspan, as the maintenance of muscle integrity is essential for preserving locomotory capacity in aging animals. The improved sarcomere organization and muscle fiber integrity observed in ADH-1^OE^ animals are consistent with the observed upregulation of small heat shock proteins, including *hsp-3*, *rpn-6.1*, *rpn-6.2*, *hsp-16.1*, *hsp-16.2*, and *hsp-60*, which are critical for maintaining muscle proteostasis(Kutzner et al., 2024; Shemesh et al., 2013; van Oosten-Hawle et al., 2013). RNAi against *hsf-1*—the master regulator of the heat-shock response—suppressed the locomotory improvements, indicating that this transcriptional axis is required for extended muscle performance in ADH-1^OE^ animals.

Intriguingly, several tissues that benefit from ADH-1 overexpression do not express the gene endogenously. We observed no detectable ADH-1 expression in GABAergic or olfactory neurons, yet ADH-1^OE^ animals maintain better axonal structure and chemotaxis later in life. This disconnect between expression and benefit supports a non-cell-autonomous mode of action, in which ADH-1 activity in muscle, pharynx, or intestine initiates systemic protective programs. Prior work has established that transcription factors such as HSF-1, DAF-16, and SKN-1 can orchestrate broad organismal responses, including cross-tissue proteostasis signaling (Prahlad et al., 2008; Taylor & Dillin, 2013c). In contrast to these transcriptional master regulators, ADH-1 is a metabolic enzyme, raising the possibility that enzymatic remodeling of metabolites and redox tone constitutes a distinct, under-appreciated mechanism of tissue crosstalk that warrants further investigation.

Rather than contrasting ADH-1 with “isolated” longevity axes, we emphasize potential mechanistic overlap with canonical pathways. In Ghaddar et al., *adh-1* activity was required for the longevity of *daf-2* mutants, *eat-2* (CR) models, and mTOR RNAi worms, suggesting that ADH-1 intersects with, or is permissive for, these pathways. This positioning is consistent with ADH-1 acting at a metabolic–proteostatic convergence point that integrates redox signaling, chaperone capacity, and inter-organ communication. Candidate distal effectors most consistent with our data include metabolites (e.g., acetate; Miskelly et al., 2019), changes in NAD⁺/NADH ratios, and redox-active small molecules. While additional mediators cannot be excluded, our current data more strongly support metabolic and redox intermediates than peptide hormones as primary drivers.

From a translational perspective, these non-cell-autonomous benefits are especially compelling. In mice Adh1, has been found enriched in EVs (Cho et al., 2017). Complementing this, in humans, ADH1C—the ortholog of *C. elegans adh-1*—has been identified as a component of the liver secretome as a plasma circulating factor (Kolker et al., 2012; Stelzer et al., 2016), and one whose circulation may be dependent on patients’ insulin sensitivity (Choi et al., 2019). This is particularly interesting for two reasons. Firstly, this suggests a potential mechanism by which ADH1 activation in one tissue could influence activity in another. Secondly, this suggests that ADH1 may play a metabolic role in humans outside of its native ethanol metabolism, potentially similar to the model proposed in Ghaddar et al. and this work via *C. elegans*.

In sum, ADH-1 overexpression confers broad protection across cellular systems by combining redox adaptation, proteostatic reinforcement, and inter-tissue exchange of beneficial effectors or signals. This integration positions ADH-1 as a regulator of healthspan architecture—capable not only of delaying decline in individual tissues but also of orchestrating organism-wide resilience. Future work should aim to map the upstream triggers of *adh-1* induction, define its metabolic and, possibly, secreted effectors, and determine whether its benefits can be mobilized pharmacologically or genetically in other species. These investigations will clarify whether ADH-1 represents a conserved axis for systemic geroprotection, and whether it can be leveraged to promote healthy aging in humans.

## Methods

### *C. elegans* Strains and Maintenance

The full list of *C. elegans* strains used in this study is provided in the Key Resources Table.

For age-related phenotype assessments, *C. elegans* were synchronized by L1 arrest: hatchlings were incubated overnight (18–24 h) in incomplete S-buffer. Approximately 200 synchronized L1s were then dispensed onto 6-cm NGM plates seeded with 200 µL of freshly harvested OP50 and incubated at 20 °C until the young adult stage. At that point, worms were transferred to fresh NGM plates supplemented with 100 µg/mL FUdR to prevent progeny production. Phenotypic assessments were performed on day 1, 2, 8, 10, and/or 12 of adulthood, depending on the assay.

### Fat Staining

Wild-type (N2) and ADH-1^OE^ (GMW20–22; see Key Resources Table) worms were grown and synchronized as described above. Oil Red O (ORO) staining was performed as previously described (Ke et al., 2018, 2021; Wählby et al., 2014). Briefly, day 1 and day 10 adults were washed four times in incomplete S-buffer, concentrated, and fixed by resuspension in 60% isopropanol. Fixed worms were stained with freshly prepared, filtered 60% ORO solution and incubated in a humid chamber at 25 °C overnight. Staining duration was empirically tested at 5, 10, and 20 h at 25 °C in a humid chamber.

Imaging was conducted on Nikon Eclipse Ti inverted microscope using a 10x Objective. Image acquisition was carried out with a DS-Ri2 Nikon camera and NIS software. Images were exported as RGB and processed in ImageJ/Fiji (fiji.sc). The blue channel was isolated; wild-type images were inverted and thresholded to capture worm bodies, and the resulting threshold parameters were then applied identically to all other conditions. Binary masks generated from this thresholding were analyzed with Analyze Particles to extract mean pixel intensity (MPI) per worm as the ORO signal. At least three independent biological replicates were performed.

### RT-qPCR

Approximately 1,500 synchronized L1 worms were seeded per 10-cm RNAi plate containing control *E. coli* XU363. Upon reaching the young adult stage, worms were transferred to control or experimental RNAi plates supplemented with 100 µg/mL FUdR. On day 8 of adulthood, animals were harvested using a mesh to remove dead eggs, flash-frozen in liquid nitrogen, and stored at −80 °C until processing.

Total RNA was extracted from frozen worms with TRI Reagent (MRC, USA) according to the manufacturer’s instructions. RNA purity and concentration were assessed by NanoDrop spectrophotometry. Three micrograms of RNA (3 µg) were used to synthesize complementary DNA (cDNA). Quantitative PCR was performed using cDNA, SYBR Green (Bio-Rad), and gene-specific primers (Table S1) on a real-time PCR thermal cycler (Bio-Rad, USA). Fold changes were calculated using the Pfaffl method (Pfaffl, 2001), and statistical significance relative to the WT control was determined using an unpaired Student’s t-test. At least three independent biological replicates were performed.

### Pharyngeal Pumping Assay

Wild-type (N2) and ADH-1^OE^ (GMW20; see Key Resources Table) worms were grown and synchronized as described in *C. elegans* Strains and Maintenance. Pharyngeal pumping was assessed on day 1 and day 8 of adulthood.

Sixty-second videos of the head region were acquired on a Zeiss Axio Zoom.V16 stereomicroscope (PlanNeoFluar Z 2.3×/0.57 FWD objective; ∼112× total magnification) equipped with a digital camera. Videos were played back at 10 fps, and pumps were counted within multiple non-overlapping 30-s windows per animal. Because animals could transiently drift out of focus during a 60-s recording, pumps per minute were calculated by averaging the counts from the available 30-s windows and extrapolating to per-minute rates. At least three independent biological replicates were performed.

### Chemoattractant assay

Wild-type (N2) and ADH-1^OE^ (GMW20; see Key Resources Table) worms were grown and synchronized as described in *C. elegans* Strains and Maintenance. Chemotaxis assays were performed on unseeded NGM plates (10 cm).

Each plate was divided into four quadrants, a 0.5-cm diameter circle was marked at the center, and a single “solution point” was designated in each quadrant, positioned equidistant from one another and ≥2 cm from the center. Solutions of attractant (diacetyl in incomplete S-buffer) and control (incomplete S-buffer) were prepared.

On day 1 and day 10 of adulthood, worms were harvested by washing plates with incomplete S-buffer, washed four times to remove residual bacteria, and concentrated. Approximately 150 worms were deposited at the center of each plate, after which control and attractant solutions were pipetted onto their respective solution points (T0). After 1hr, assays were analyzed following the protocol detailed in (Margie et al., 2013). At least three independent biological replicates were performed.

### Paraquat Challenge

Wild-type (N2) and ADH-1^OE^ (GMW20; see Key Resources Table) worms were grown and synchronized as described in *C. elegans* Strains and Maintenance. Oxidative stress assays were performed as previously described (Mony et al., 2021; Senchuk et al., 2017)with minor modifications. Six-centimeter NGM plates were seeded with fresh OP50. Paraquat (methyl viologen) was added to achieve a final plate concentration of 100mM; plates were placed on an orbital shaker for 1 h to ensure even distribution and then dried in a biosafety hood for 20 min.

Day 1 and day 8 adults were transferred to paraquat plates by picking. Unless otherwise noted, ∼20–30 worms were placed per plate, with ≥3 plates per condition (experimental replicates), and incubated at 20 °C. Survival was scored at regular intervals until at least 50% mortality was reached. Animals that crawled off the agar, ruptured, or displayed internal hatching were censored. At least three independent biological replicates were performed.

### Thrashing Assay

Wild-type (N2) and ADH-1^OE^ (GMW20–22; see Key Resources Table) worms were grown and synchronized as described in *C. elegans* Strains and Maintenance. Thrashing was assessed in day 1 and day 12 adults.

Twenty-four–well plates were pre-filled with 2 mL S-buffer per well. Using a wire pick, individual worms were placed into separate wells directly into the buffer. After a 2-4 min acclimation (kept ≤5 min uniformly across animals), 60-s videos of each well were acquired sequentially on a ZEISS Axio Zoom.V16 stereomicroscope (PlanNeoFluar Z 2.3×/0.57 FWD objective; ∼112× total magnification) using ZEISS ZEN software (RRID:SCR_013672). Movies were analyzed in ImageJ/Fiji (RRID:SCR_002285) (Schindelin et al., 2012) and the WrmTrck plugin(Nussbaum-Krammer et al., 2015) to quantify body bends per second (BBPS). A “body bend/thrash” was defined as one complete change in lateral bending of the mid-body (left→right→left or right→left→right). When acquisition frame rates differed between runs, bend counts were normalized to elapsed time to report BBPS. At least three independent biological replicates were performed.

### Fluorescent Imaging and Processing

Imaging was performed on either (1) a Nikon Eclipse Ti inverted microscope equipped with a Yokogawa CSU-X1 spinning disk, a Hamamatsu ORCA-Flash 4.0 sCMOS camera, and 405/488/561/640-nm lasers using a Plan Apo 10x, or (2) a Leica Stellaris 8 Fluorescent Spinning Disk Confocal with a 63×/1.4 NA oil-immersion objective. GABAergic motor neurons and body-wall muscles were imaged on the Leica system. PolyQ aggregates, reference images of ADH-1 tissue localization, HyPer::YFP images, and *hsp-4*p::GFP images were acquired on the Nikon system.

For all imaging, worms were immobilized in 10 mM levamisole on 2% agarose pads.. For experiments requiring fluorescent imaging, Z-stacks were collected. Equal laser intensity and exposure time was used for each replicate of a respective experiment. Image processing was solely conducted in ImageJ/Fiji (fiji.sc) software. Maximum projections were generated in almost all cases. In some cases, excessive worm movements made maximum projections not possible. In those instances, the brightest single slices were chosen from each respective group. Images then had rolling background subtraction applied (radius of 50) except in cases where thresholding was used (eg. PolyQ imaging). Per worm data was collected via manual worm tracing in Fiji. For *hsp-4*::GFP and HyperROS imaging, worm ROIs were drawn from the first two intestinal cells to the tails of the animal. This was done to minimize any false positive signal from the green pharynxes of ADH-1^OE^ animals. Animals were excluded if they were determined to be much younger than the majority of worms in the image (eg. smaller total area discrepancy and/or lack of eggs). Due to the green pharynx of ADH-1^OE^ worms, blinding was not possible in most cases. Blinding was conducted in the muscle scoring analysis as muscle images were taken from worm midbodies (pharynx not visible) and the scoring system was more subjective so blinding was heavily prioritized. Then, properties of each ROI were measured using the Fiji “measure” function. Data analyses was then conducted in R (see Statistical Testing).

### Muscle Integrity Assay

RW1596 and GMW031 (see Key Resources Table) *C. elegans* were grown and synchronized as described in *C. elegans* Strains and Maintenance. The mid-body region was imaged to avoid distortions near the vulva, head, and tail. Z-stacks were acquired on a Leica Stellaris 8 fluorescent confocal microscope using a 40× objective with 2 µm optical steps. Images were collected at days 1, 6, 8, and 10; data shown are from day-10 adults. Image acquisition and processing followed the workflow described in “Fluorescent Imaging and Processing”. Images were blinded using Blinder (Cothren et al., 2018) prior to scoring, then categorized into three classes: healthy, intermediately damaged, or severely damaged as exemplified in DataS4E-G. Two independent biological replicates were performed.

### Neuronal integrity Assay

CZ13799 and GMW028 (see Key Resources Table) worms were grown and synchronized as described in *C. elegans* Strains and Maintenance. Animals were immobilized in 10 mM levamisole on 2% agarose pads, and VD/DD GABAergic axonal commissures were imaged from head to tail on a Leica Stellaris 8 fluorescent spinning-disk confocal microscope using a 63×/1.4 NA oil-immersion objective. Z-stacks were acquired at 1 µm optical steps. For image processing, the workflow described in “Fluorescent Imaging and Processing” was followed. Specific to this assay, commissural axons were cropped away from surrounding tissue to prevent spurious detections. Neurodegeneration was quantified in ImageJ/Fiji (fiji.sc) using the FindFoci plugin (Herbert et al., 2014). Axonal “stippling” events were operationally defined as punctate GFP spikes above the commissural background. FindFoci was initialized by establishing a baseline detection threshold on a manually curated training set of healthy commissures; this threshold and all analysis parameters were then fixed and applied uniformly to all images and conditions. For each worm, spike counts were computed per commissure and then averaged across all commissures to yield a single value (mean stippling events per commissure per worm). At least two independent biological replicates were performed.

### Polyglutamine aggregate assessment

AM140 and GMW0024(see Key Resources Table) worms were grown and synchronized as described in *C. elegans* Strains and Maintenance. For imaging, animals were immobilized in 1 mM levamisole on 2% agarose pads. Z-stacks were acquired at 0.9 µm optical steps with a 20× objective on the Nikon spinning-disk system (Eclipse Ti + Yokogawa CSU-X1; Hamamatsu ORCA-Flash 4.0). Whole-animals were tiled and stitched in NIS-Elements (RRID:SCR_014329) using linear blending with 50% overlap.

Image analysis was performed in ImageJ/Fiji (RRID:SCR_002285)(Schindelin et al., 2012). Worm outlines were cropped to generate per-animal ROIs. Aggregates were enhanced for detection by local automatic thresholding (Otsu, default settings) benchmarked against WT animals and then applied to ADH-1^OE^ animals. The same threshold was applied to all images within a replicate. Particles were then segmented with Analyze Particles (size: 2–500; other parameters at default). This size window was chosen to exclude GFP-positive pharynxes in ADH-1^OE^ animals and to avoid counting large diffuse GFP signals as large aggregates. All processing parameters were applied identically across conditions. Two independent biological replicates were performed.

### Statistical Testing

All analyses and visualizations were performed in R (R Foundation) using RStudio (Posit). Core packages included: Tidyverse(*Tidyverse*, n.d.), ggplot2(*Authors and Citation*, n.d.), ggpubr(*Ggplot2 Based Publication Ready Plots*, n.d.), patchwork(*Getting Started*, n.d.), RColorBrewer(Neuwirth, 2022), survival(Therneau et al., 2024), ggsurvfit(Sjoberg et al., 2024), dplyr(Wickham, François, et al., 2023), readxl(Wickham et al., 2025), infer(Bray et al., 2025), svglite(Wickham, Henry, et al., 2023), survminer(Kassambara et al., 2024), glue(Hester et al., 2024), ggtext(Wilke & Wiernik(@bmwiernik), 2022), rstatix(Kassambara, 2023), knitr(Xie [aut et al., 2025), stringr(Wickham, Software, et al., 2023), and extrafont(Chang, 2023). Data were assessed for normality (Shapiro–Wilk test). When assumptions were met, two-sided t-tests (Welch’s correction when variances were unequal). When assumptions were violated, Wilcoxon/Mann–Whitney or Kruskal–Wallis tests were applied with appropriate multiple-comparison procedures. Survival data were analyzed by Kaplan–Meier estimates and log-rank tests. The specific test used for each comparison is stated in the corresponding figure legend.

