## Supplemental Figures for "Broad Effects of Activation of Alcohol Dehydrogenase 1 on Healthspan Extension"

### Supplementary Data

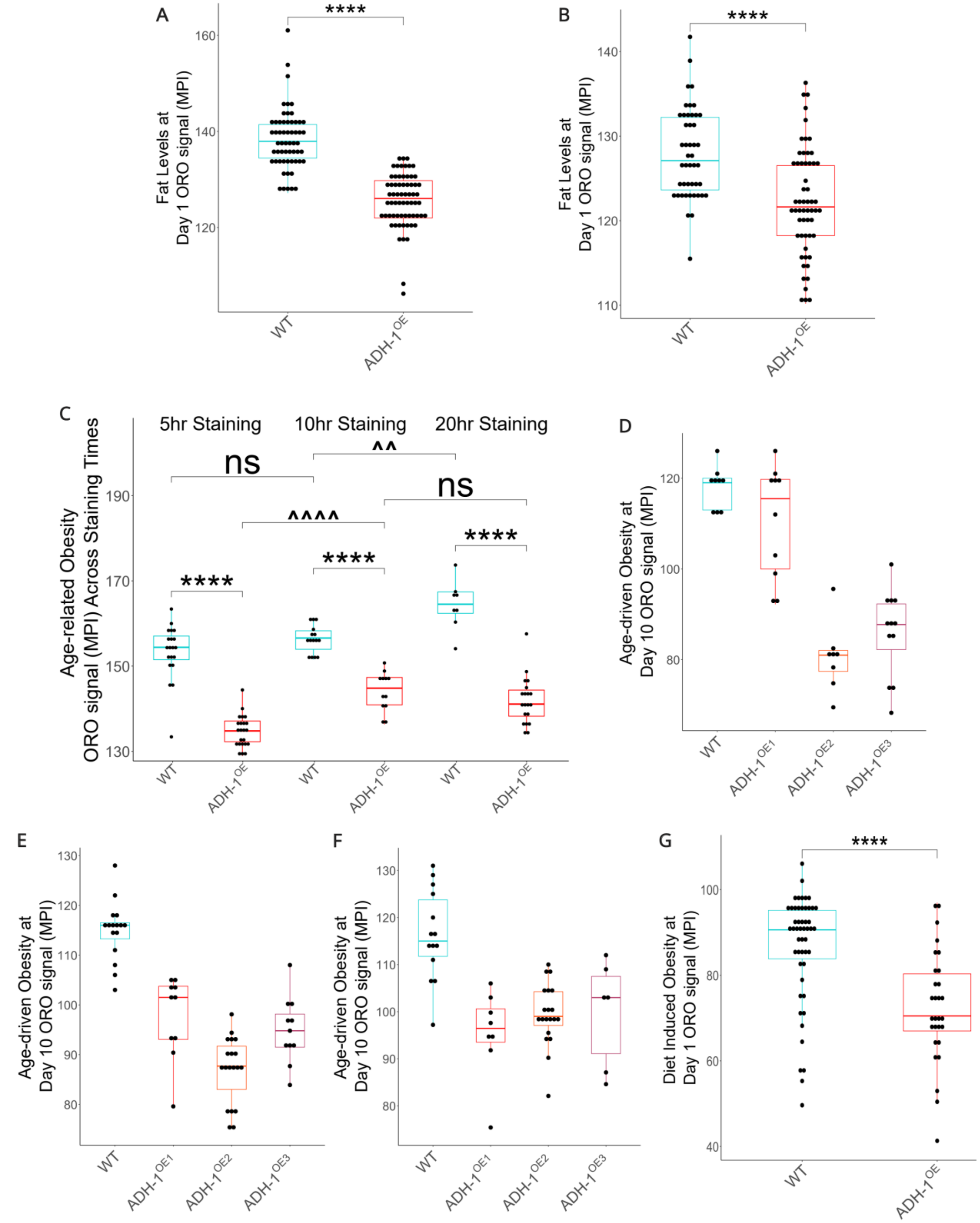

**DataS1**: (A & B) Two independent biological replicates showing that ADH-$1^{OE}$ adults are leaner then WT animals, (C) Worms were stained with oil redO (ORO) for 3 different lengths of time, all other variables were maintained equal. As denoted with asterisks, staining time does not determine the difference in ORO Mean Pixel Intensity (MPI) between ADH-$1^{OE}$ and WT aging *C. elegans* (8 days of adulthood). Nevertheless, as denoted with carets (^), longer staining times did increase the ORO signal. ADH-$1^{OE}$ ORO uptake seems to plateau between 10 and 20h, (D-F) Each panel depicts an independent biological replicate showing that three independent ADH-$1^{OE}$ transgenic lines are resistant to age-driven obesity (day 10 adulthood), (G) Representative biological replicate showing that day 1 ADH-$1^{OE}$ animals remain leaner than WT when fed an obesogenic diet (55mM Fructose). Statistical testing was conducted using Welch's t-test after outlier correction or Wilcoxon when populations were non-normal (Panels D, E, F, and G), Statistical testing was conducted using Welch's t-test after outlier correction. Error bars denote SEM. ns= not significant, *$p<0.05$, **$p<0.01$, ***$p<0.001$, ****$p<0.0001$, **MPI** = Mean Pixel Intensity.

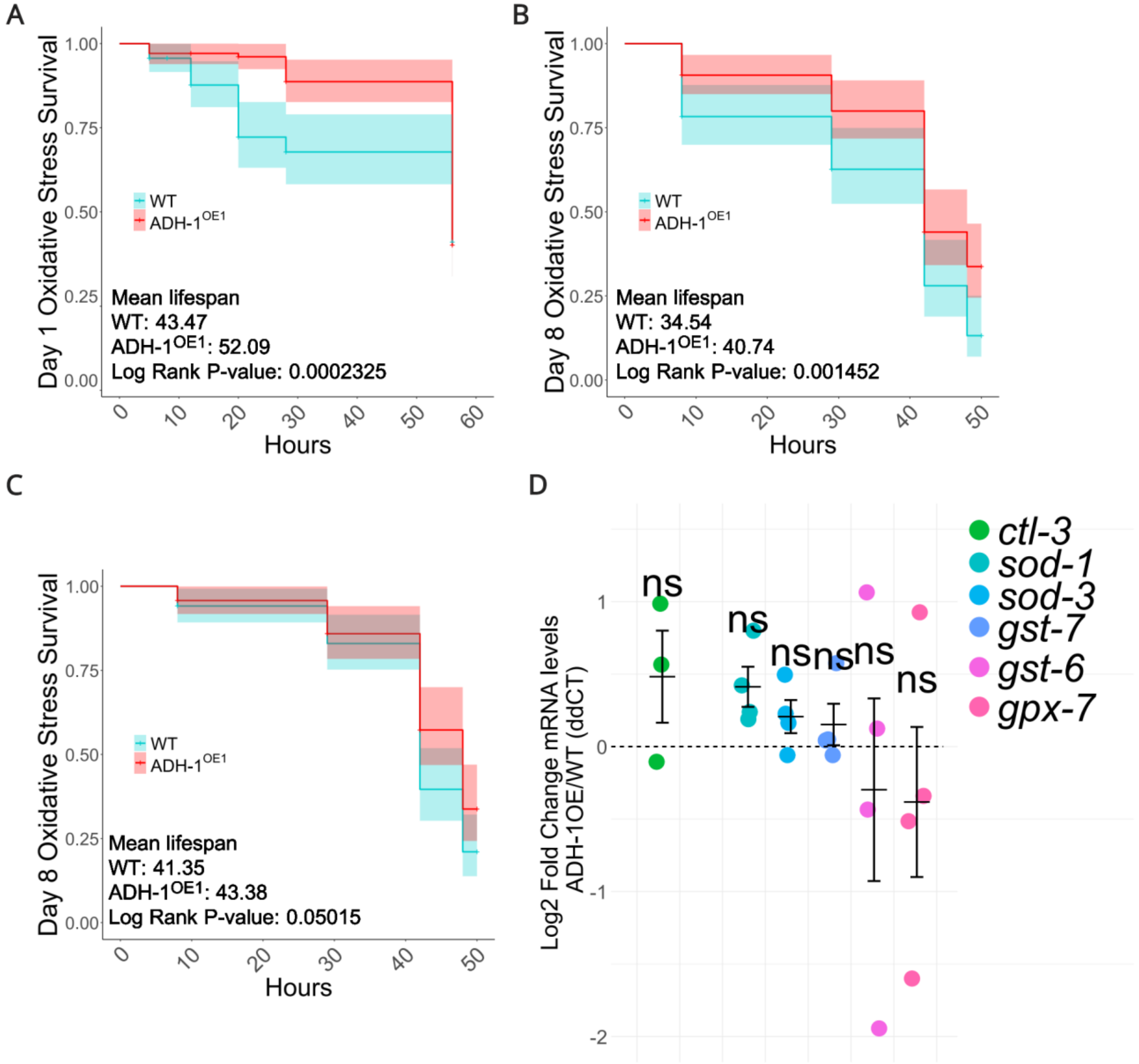


**DataS2:** (A) An independent trial showing ADH-1$^{OE}$ animals survive better than WT to oxidative stress exposure at day 1, (B) An independent trial showing ADH-1$^{OE}$ animals survive better than WT to oxidative stress exposure at day 8, (C) An independent trial showing ADH-1$^{OE}$ animals survive better than WT to oxidative stress exposure at day 8, (D) Redox genes tested by qPCR that did not show significance despite some trends, Statistical testing was conducted using Welch's t-test after outlier correction. Statistical testing of survival conducted by Kaplan-Meier Log-Rank Fold test. Error bars denote SEM. ns= not significant, *p<0.05, **p<0.01, ***p<0.001, ****p<0.000

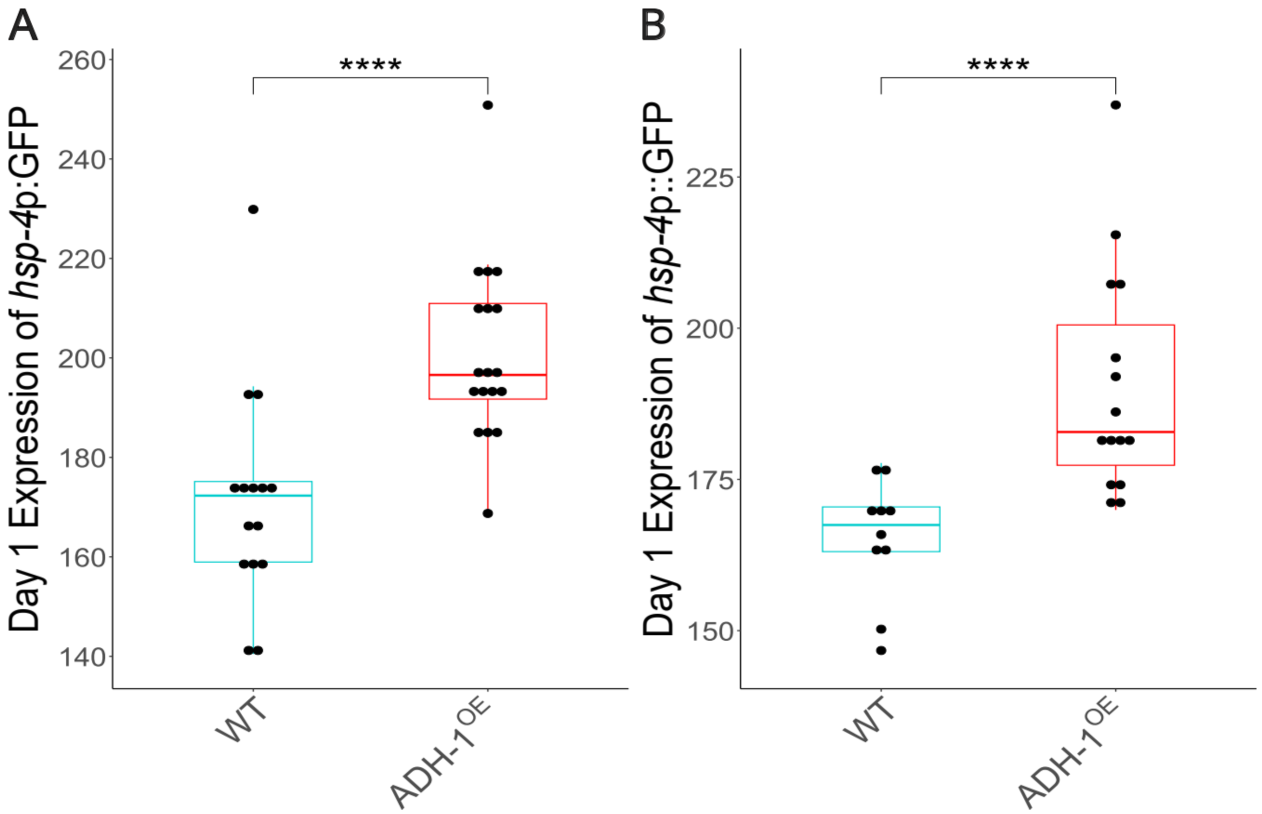


**DataS3:** (A) An independent trial showing ADH-1$^{OE}$ animals have higher levels of hsp-4 expression via *hsp-4*p::GFP, (B) An independent trial showing ADH-1$^{OE}$ animals have higher levels of hsp-4 expression via *hsp-4*p::GFP. Statistical testing was conducted using Wilcoxon due to having a nonnormal population after outlier correction. Error bars denote SEM. ns= not significant, *p<0.05, **p<0.01, ***p<0.001, ****p<0.0001.

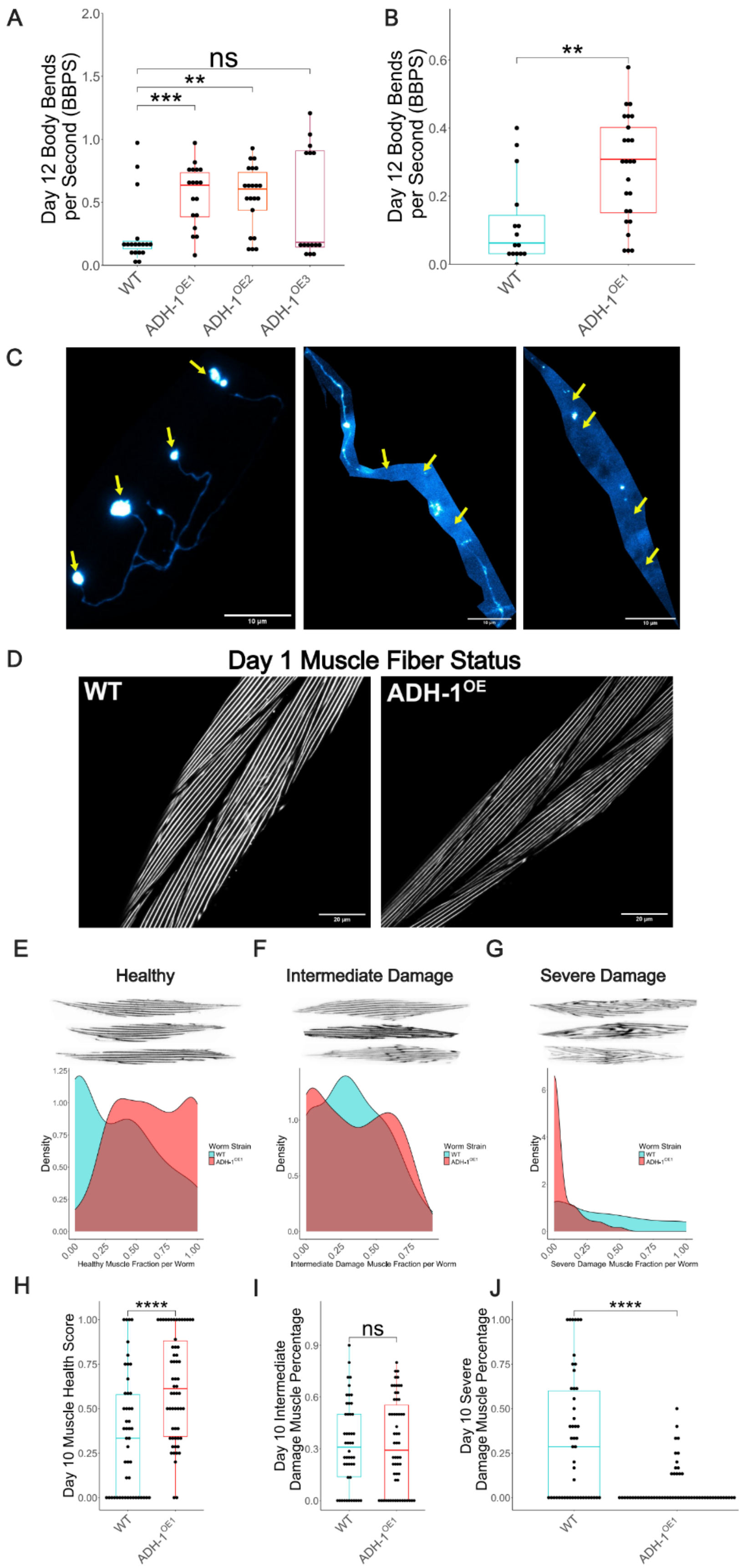
A
Day 12 Body Bends per Second (BBPS)
ns
**
***
WT
ADH-1OE1
ADH-1OE2
ADH-1OE3
B
Day 12 Body Bends per Second (BBPS)
**
WT
ADH-1OE1
C
D
Day 1 Muscle Fiber Status
WT
ADH-1OE
E
Healthy
F
Intermediate Damage
G
Severe Damage
Density
Worm Strain
WT
ADH-1OE1
Healthy Muscle Fraction per Worm
Intermediate Damage Muscle Fraction per Worm
Severe Damage Muscle Fraction per Worm
H
Day 10 Muscle Health Score
****
I
Day 10 Intermediate Damage Muscle Percentage
ns
J
Day 10 Severe Damage Muscle Percentage
****

**DataS4:** (A) An independent trial showing three independently generated ADH-$1^{OE}$ strains outperform WT in thrashing at day 12, (B) An independent trial showing the ADH-$1^{OE}$ strain outperforms WT in thrashing at day 12, (C) Three example images of WT GABAergic neuron commissures as observed in day-12 WT adults, (D) At day 1, WT and ADH-$1^{OE}$ animals display similarly well-structured sarcomeres, (E-G) Density plots of healthy, intermediate damage, and severe damage muscle fibers. (H-J) Corresponding boxplot quantifications of said density plots showing that while healthy and severely damaged muscle fibers differed significantly between ADH-$1^{OE}$ and WT animals, intermediately damaged muscle fibers showed no difference. Statistical testing was conducted using Welch's t-test after outlier correction or Wilcoxon when populations were non-normal (Panels A, B, G, H, and I). Error bars denote SEM. ns= not significant, *p<0.05, **p<0.01, ***p<0.001, ****p<0.0001, **MPI** = Mean Pixel Intensity.

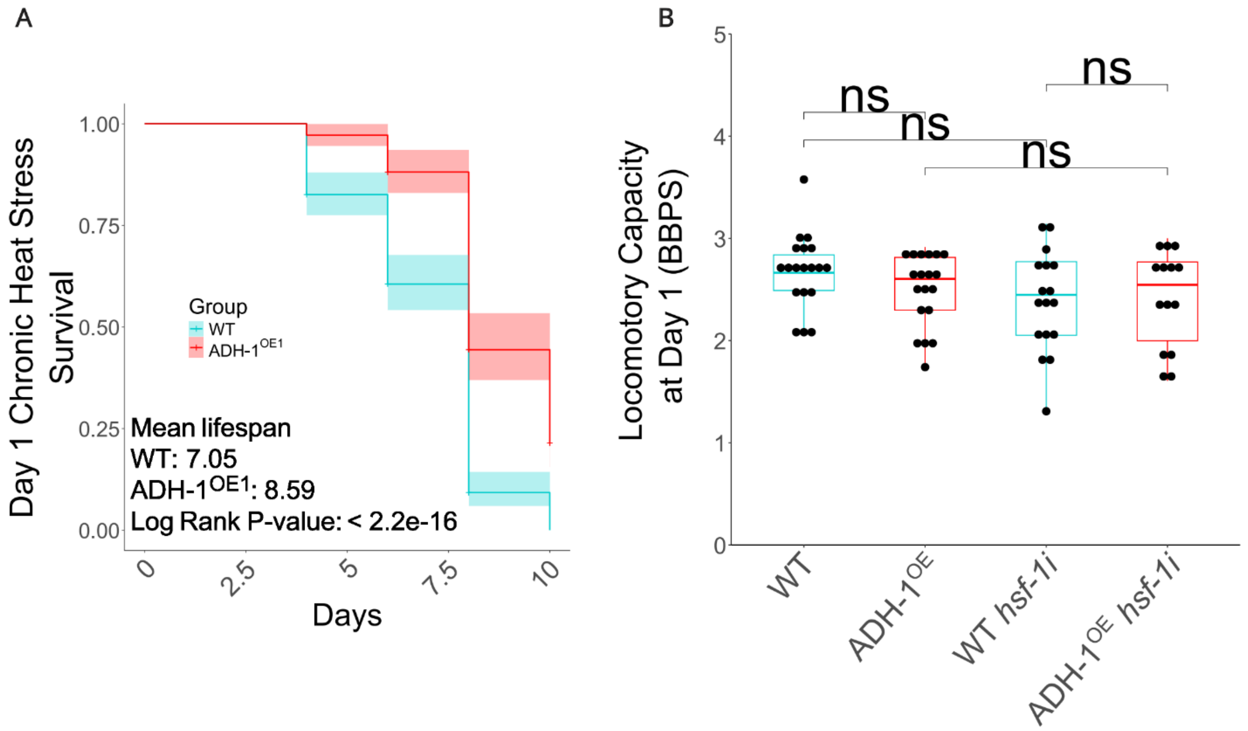


**DataS5:** (A) Representative survival curve showing that ADH-1$^{OE}$ animals better survive chronic heat stress, (B) *hsf-1i* has no impact on locomotory capacity at day 1 (biological replicate=1). Statistical testing was conducted using Welch's t-test after outlier correction and, for survival, by Kaplan-Meier Log-Rank Fold test. Error bars denote SEM. ns= not significant, *p<0.05, **p<0.01, ***p<0.001, ****p<0.0001, **MPI** = Mean Pixel Intensity.
